# Whole-genome sequencing of ethnolinguistic diverse northwestern Chinese Hexi Corridor people from the 10K_CPGDP project suggested the differentiated East-West genetic admixture along the Silk Road and their biological adaptations

**DOI:** 10.1101/2023.02.26.530053

**Authors:** Guanglin He, Hongbing Yao, Qiuxia Sun, Shuhan Duan, Renkuan Tang, Jing Chen, Zhiyong Wang, Yuntao Sun, Xiangping Li, Shaomei Wang, Liping Hu, Libing Yun, Junbao Yang, Jiangwei Yan, Shengjie Nie, Yanfeng Zhu, Chao Liu, Mengge Wang

## Abstract

The ancient Silk Road served as the main connection between East and West Eurasia for several centuries. At any rate, the genetic exchange between populations along the ancient Silk Road was likely to leave traces on the contemporary gene pool of local people in Northwest China, which was the passage of the Northern Silk Road. However, genetic sources from northwestern China are under-represented in the current population-scale genomic database. To characterize the genetic architecture and adaptative history of the Northern Silk Road ethnic populations, we performed whole-genome sequencing on 126 individuals from six ethnolinguistic groups (Tibeto-Burman (TB)-speaking Tibetan, Mongolic (MG)-speaking Dongxiang/Tu/eastern Yugur, and Turkic (TK)-speaking Salar/western Yugur) living in Gansu and Qinghai in the 10K Chinese people Genomic Diversity Project (10K_CPGDP). We observed ethnicity-related differentiated population structures among these geographically close Northwest Chinese populations, that is, Salar and Tu people showed a close affinity with southwestern TB groups, and other studied populations shared more alleles with MG and Tungusic groups. Overall, the patterns of genetic clustering were not consistent with linguistic classifications. We estimated that Dongxiang, Tibetan, and Yugur people inherited more than 10% West Eurasian ancestry, much higher than that of Salar and Tu people (<7%). Hence, the difference in the proportion of West Eurasian ancestry has primarily contributed to the genetic divergence of geographically close Northwest Chinese populations. The signatures of natural selection were identified in genes associated with cardiovascular system diseases or lipid metabolism related to triglyceride levels (e.g., *PRIM2, PDE4DIP, NOTCH2, DDAH1, GALNT2*, and *MLIP*) and developmental and neurogenetic diseases (e.g., *NBPFs 8/9/20/25P*, etc.). Moreover, the *EPAS1* gene, a transcription factor regulating hypoxia response, showed relatively high PBS values in our studied groups. The sex-biased admixture history, in which the West Eurasian ancestry was introduced primarily by males, was identified in Dongxiang, Tibetan, and Yugur populations. We determined that the eastern-western admixture occurred ∼783–1131 years ago, coinciding with the intensive economic and cultural exchanges during the historic Trans-Eurasian cultural exchange era.

## INTRODUCTION

Archaeological, linguistic, anthropological, paleogenomic, and genetic studies have provided sufficient evidence to demonstrate the multiple migration and admixture events between the ancestors of anatomically modern humans (Lazaridis, et al. 2014; Nielsen, et al. 2017; GenomeAsia 2019; Bergstrom, et al. 2020a; Liu, Mao, et al. 2021). Archaic and ancient human activities (migration, admixture, replacement and introgression) have resulted in population separations followed by isolation, repeated population bottlenecks, and subsequent intermixing, consequently shaping modern human populations’ complex patterns of genetic diversity (Reich 2018). Human genetic studies from the expanded 1000 Genomes Project (1KGP), Human Genetic Diversity Project (HGDP), and other regional genomic projects have advanced our understanding of human genetic discovery and corresponding biological and phenotypic consequences of the genetic variations (Bergstrom, et al. 2020b; Collins, et al. 2020; Taliun, et al. 2021; Byrska-Bishop, et al. 2022; Halldorsson, et al. 2022). Genetic scientists consistently suggested that high-depth sequenced human genomes from ethnolinguistically and geographically diverse ethnic groups were necessary to capture a complete picture of human genomic diversity (Sirugo, et al. 2019). Recent large-scale genomic projects mainly focused on Europeans, which may cause European bias in human genetic research and introduce health inequality in the precision medicine era, such as the Genome Aggregation Database (gnomAD) (Collins, et al. 2020), Trans-Omics for Precision Medicine (TOPMed) program (Taliun, et al. 2021) and UK Biobank (Halldorsson, et al. 2022). However, high-coverage whole-genome sequencing studies in East Asian populations have so far mainly focused on demographically larger Han populations (Jeon, et al. 2020; Ma, Yang, et al. 2021; Zhang, Luo, et al. 2021) or been targeted at specific metabolic diseases (Cao, et al. 2020). The underrepresentation of ethnolinguistically diverse East Asians impedes our complete understanding of modern humans’ evolutionary history, genetic discovery, and human health. To comprehensively capture the global genomic diversity of under-studied Chinese ethnolinguistically diverse populations, we launched the 10K Chinese people Genomic Diversity Project (10K_CPGDP), which aimed to illuminate the population formation, biological adaptation and medical relevance of anthropologically-informed Chinese populations from rural countries via whole-genome sequencing. The first phase successfully whole-genome-sequenced 2000 genomes from more than 30 ethnolinguistically diverse populations. One of the programs of this project focused on the genetic admixture landscape and adaptative history of Northwest Chinese populations.

Northwest China, including the provinces of Gansu, Qinghai, and Shaanxi and the autonomous regions of Xinjiang and Ningxia, borders the countries of Central Asia, South Asia, Russia, and Mongolia and is located at the junction between the Eurasian steppes and the Central Plains of China. This area, now widely inhabited by Sinitic, Tibeto-Burman (TB), Mongolic (MG), and Turkic (TK)-speaking ethnic groups, is a strategic location for genetic and cross-cultural communications between West Eurasia and inland China in prehistoric and historical periods (Xu, et al. 2008; Feng, et al. 2017; Ning, et al. 2019; Ma, Yang, et al. 2021). Additionally, Northwest China connected lowland East Asia and the highland Tibetan Plateau and served as the vital buffer area for ancient people to settle on the Qinghai-Tibet Plateau (Ding, et al. 2020). Hexi Corridor in Northwest China was the critical geographical passway in the ancient Silk Road and played a pivotal function in the globalization of ancient millet, barley agriculture, and human populations. This historically documented crucial trade route facilitated human migrations and cultural exchanges between East and West Eurasia, resulting in genetic admixture between originally isolated populations and shaping the human diversity of contemporary Eurasians (Feng, et al. 2017; Li, Zou, et al. 2020; Ma, Yang, et al. 2021). The human genetic survey in this region mainly focused on the Hui and Uyghur people and ancient populations at the western end of this corridor. Xu and Pan et al. found that TK-speaking Uyghur has more western Eurasian-related ancestry than Hui and Han Chinese and harbored ancestry from South Asian, European, Siberian and East Asian populations (Pan, et al. 2022). Kumar et al. reported the largest-scale ancient genomic study, including 201 individuals from 39 archaeological sites in Xinjiang, and found that East and West Eurasian populations have been influencing different regions of Xinjiang from the Bronze Age to the historical period (Kumar, et al. 2022). Zhang et al. recently identified that the Early and Middle Bronze Age Tarim individuals have arisen from a genetically isolated local origin. The Early Bronze Age Dzungarian individuals primarily exhibited Afanasievo ancestry with an additional local contribution (Zhang, Ning, et al. 2021). Ning et al. also illustrated that the Iron Age people from Xinjiang were highly structured with East and West Eurasian ancestries (Ning, et al. 2019). Most East Eurasian ancestry was related to Northeast Asian populations, and the West Eurasian ancestry was best represented by Yamnaya-like lineage. Complex patterns of ancient population interactions and admixture occurred among ancient northwestern Chinese populations. However, to what extent West Eurasian ancestry contributed to the gene pool of modern Hexi Corridor populations remains unclear.

Northwest China is characterized by exceptionally high levels of ethnic diversity, which possesses many ethnolinguistically diverse populations, including Han Chinese and other significant ethnic minorities such as TK-speaking Uyghur, Kazakh, Kyrgyz, Salar, Tajik, Evenki, Russian, Uzbek and Tatar, MG-speaking Mongolian, Dongxiang, Tu, Xibe, Bonan, Yugur, Oroqen and Daur, TB-speaking Tibetan, and other Hui, Manchu (Zang 2016). Previous genetic studies of demographically diverse populations from Northwest China were primarily based on forensically relevant genetic markers (Yao, et al. 2012; Wei, et al. 2014; Meng, et al. 2015; Yao, et al. 2016; Wang, et al. 2018; Zhan, et al. 2018). Most of these studies aimed to evaluate the forensic efficacy and allele frequency spectrum of those markers in Northwest Chinese populations. Recent studies based on the genome-wide SNP data have revealed that present-day people in this region exhibited an apparent genetic admixture of East Asian and West Eurasian ancestries (Feng, et al. 2017; Zhao, et al. 2020; Ma, Chen, et al. 2021; Ma, Yang, et al. 2021; Yao, et al. 2021; Pan, et al. 2022). Feng et al. identified the southwest and northeast differentiation between Xinjiang Uyghurs and described the genetic makeup of Uyghurs as an admixture of approximately 25-37% European, 12–20% South Asian, 15–17% Siberian, and 29–47% East Asian ancestries (Feng, et al. 2017). Zhao et al. found that Xinjiang Mongolians and Kazakhs derived about 6– 40% of their ancestry from West Eurasians (Zhao, et al. 2020). Yao et al. identified the signature of West Eurasian admixture in northwestern Hans, dating back to around 1000 CE (Yao, et al. 2021). Ma et al. conducted one multiple population-based work and estimated that Qinghai minorities inherited 9–15% West Eurasian ancestry and identified the sex-biased admixture in these ethnic groups from Northwest China (Ma, Chen, et al. 2021). However, array-based genetic studies missed some rare or previously unknown genetic variations, which may provide more information for biological adaptation and cause bias in admixture modeling (Bergstrom, et al. 2020a). Recently, Ma et al. whole-genome-sequenced the Ningxia Hui people and dissected the ancestry compositions of Huis living in Ningxia at a fine scale (Ma, Yang, et al. 2021). They found that Ningxia Huis derived less West Eurasian ancestry (∼10%) than Uyghurs. However, it is remarkable that most whole-genome-based studies have been targeted at demographically sizeable ethnic groups. Fully discovering and understanding the full spectrum of variations via whole-genome sequencing studies in Northwest China is limited. The underrepresentation of northwestern ethnic groups with relatively small population sizes via whole-genome studies has limited the dissection of the peopling history of Northwest China, presentation of the full spectrum of genetic variations and the illumination of the medical relevance and biological adaptation.

In this study, we performed whole-genome sequencing on 126 Northwest Chinese individuals belonging to five ethnic groups from three language families (Yugur, Dongxiang, Tu, Salar, and Tibetan) to characterize the genomic diversity comprehensively, infer the admixture history and characterize the biological adaptative features of Northwest Chinese populations. The Yugurs are a TK/MG-speaking ethnic group living primarily in Sunan Yugur Autonomous County in Gansu. The TK-speaking Yugurs mainly live in the West of the County, and the MG-speaking Yugurs mainly live in the County’s eastern part. The Dongxiang people are an MG ethnic group primarily residing in Linxia Hui Autonomous Prefecture and surrounding areas of Gansu in Northwest China. The paternal profile showed that most Dongxiang people belonged to haplogroup R1a1a-M17 (Shou, et al. 2010). The Tu people are also an MG ethnic group mainly living in Qinghai and Gansu provinces. Xu et al. found that haplogroups R1a1a-M17, D1a1a-M15, O2a2b1a1-M117, and O2a1b-IMS-JST002611 were the most frequent paternal lineages in Tu people (Xu and Wen 2017). The Salar people live primarily in the Gansu-Qinghai border region and are a TK ethnic minority. The paternal genetic lineages of Salar people exhibited a mix of East Asian and West Eurasian haplogroups, while their maternal lineages were overwhelmingly of East Asian origin (Wang, et al. 2003; Ma, Chen, et al. 2021). The Tibetan people are a TB-speaking ethnic group native to Tibet Autonomous Region, in addition, significant numbers of Tibetans reside in Gansu, Qinghai, Sichuan, and Yunnan provinces, as well as in Nepal, India, and Bhutan. Tibetans living in Gansu and Qinghai speak Amdo Tibetan, which is one of the three branches of traditional classification (the other two being ÜTsang and Khams Tibetan) of Tibetic languages (Gelek2017). Here, we analyzed the allele and haplotype sharing within Northwest Chinese groups and compared them with modern and ancient Eurasian populations. This will provide new insights into the complex admixture history of Northwest Chinese populations and advance the understanding of intensive contacts between East and West Eurasians.

## RESULTS

### Data quality and novel variants

We sequenced 126 self-identified ethnolinguistically indigenous people in Northwest China at a mean depth of 13.20X. Genotype calling and quality controls were carried out jointly with 126 sequenced samples. The final dataset consisted of 16,357,343 single nucleotide polymorphisms (SNPs) and 1,899,000 insertions or deletions (InDels) from 22 autosomes and the X chromosome. In our call set, 4,694,877 SNPs (28.7%) and 721,808 InDels (38.0%) were not cataloged in the dbSNP 138 version. Notably, we discovered 15,409,525 bi-allelic autosomal SNPs; singletons accounted for 30.2% of all the bi-allelic autosomal SNPs, 41.1% reached minor allele frequency (MAF) < 0.01, and 19.2% had 0.01 ≤MAF < 0.05. Unsurprisingly, most novel variants were extremely rare, with singletons accounting for 68.5% of the novel variants, 82.2% of the novel variants reaching MAF < 0.01, 13.5% reaching 0.01 ≤MAF < 0.05, and only 4.3% of the novel variants being common. We estimated the transition versus transversion ratio (Ti/Tv) of 2.04 across 15.4 million bi-allelic autosomal SNPs, which was consistent with the values reported by previous whole-genome studies (Consortium, et al. 2015; Genomes Project, et al. 2015; Wu, et al. 2019).

### Overview of genetic affinity and population structure

To investigate the general patterns of relatedness between studied Northwest Chinese populations and Eurasian populations, we began with a principal component analysis (PCA) by analyzing the whole-genome sequencing dataset panel. PCA results in the context of Eurasian populations showed that target populations sat between East Asian and West Eurasian clusters but were more closely related to East Asian people (**supplementary fig. S1A**). Notably, though target Northwest Chinese populations were placed together with East Asians, they were genetically distinguishable from linguistically close populations and deviated toward western Eurasians (**supplementary fig. S1B**). We then dissected the genetic relationship between target people and Eurasian populations at much larger temporal and spatial scales by merging our data with modern and ancient Eurasian populations from the Human Origins (HO) dataset and conducted a PCA analysis in the context of Eurasians (Lazaridis, et al. 2014). Northwest Chinese populations were located on the northern branch of the north-south cline across East Asia; these sampled individuals largely overlapped with TB and MG groups from North China (**supplementary fig. S2**). Besides, they also exhibited a close relationship with ancient populations from Nepal, Yellow River Basin (YRB), West Liao River (WLR, represented by Late Neolithic to Bronze Age ancients), Mongolia (represented by Late Medieval Mongols), and Japan (**fig. 1A and supplementary fig. S2**). We further explored the genetic coordinates of Northwest Chinese populations in the context of East Asia (**supplementary fig. S3**). In general, target populations had a close relationship with YRB farmers from the Central Plain. Specifically, Dongxiang, Yugur, and Tibetan populations were grouped with MG-speaking groups from North China, while Salar and Tu populations clustered together with TB groups and ancient populations from Nepal, Upper YRB, Inner Mongolia, and Shaanxi as well as Coastal Early Neolithic northern East Asians. Notably, an enlarged view of the northern East Asian coordinates revealed a trend of Zhuoni Tibetan tilted toward reference northwestern TB-speaking groups, while studied Salar and Tu populations largely overlapped with southwestern TB-speaking references (**fig. 1B**).

**Figure 1.**
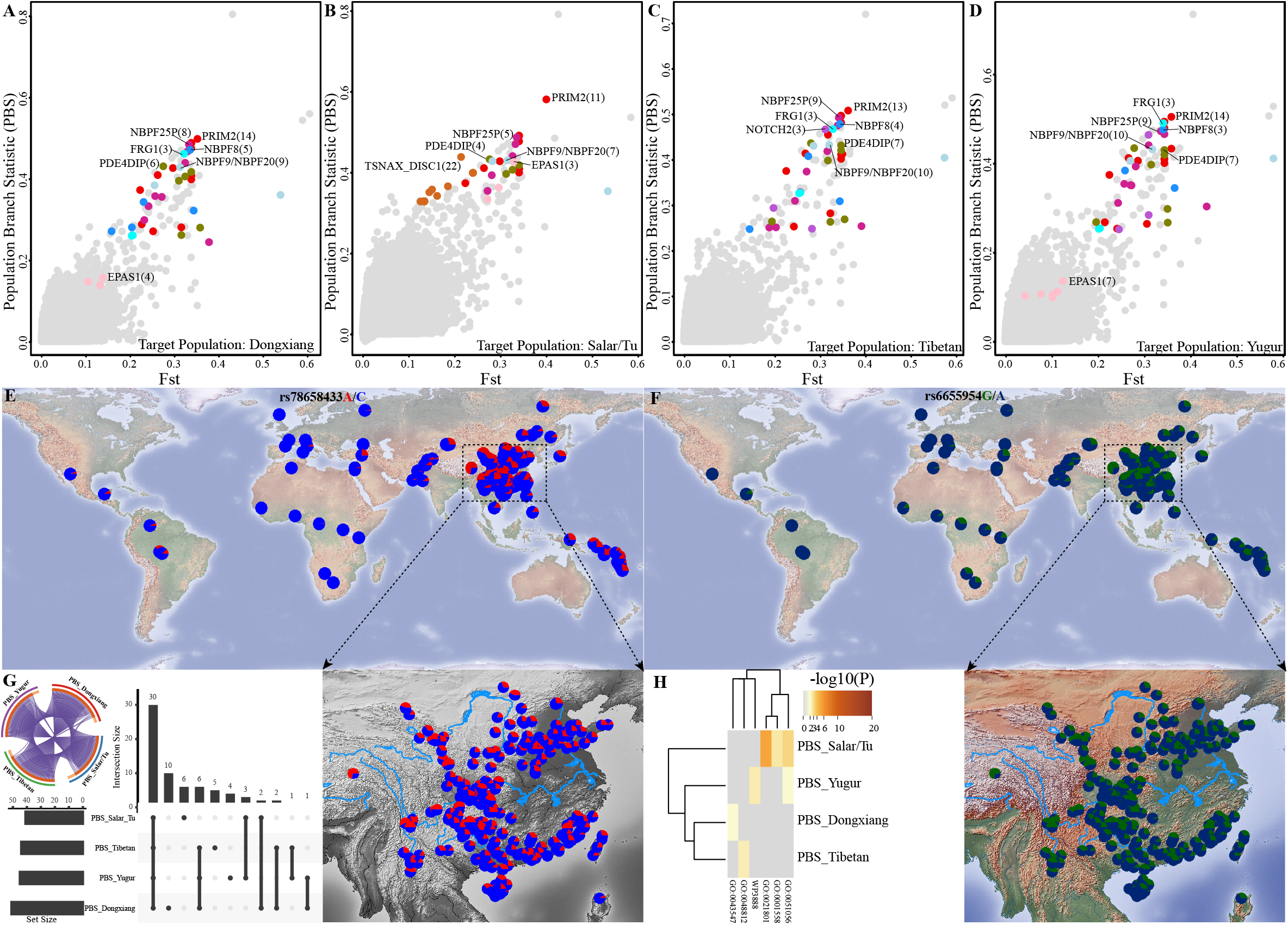
Population structure of Northwest Chinese populations and modern/ancient Eurasians based on the merged HO dataset. (**A**), Principal component analysis (PCA) of 373 modern and ancient Eurasian populations, in which modern Mlabri and ancient populations were projected onto the first two PCs. Modern reference populations were color-coded gray, and clusters or clines of ancient populations are labeled on the plot. (**B**), Patterns of genetic relationship between target Northwest Chinese populations and 82 modern and ancient northern East Asian populations. (**C**), The results of outgroup *f*_*3*_-statistics of the form *f*_*3*_(Target, Modern Eurasians; Mbuti). MG, Tungusic, TK, and TB groups were employed as modern Eurasian reference populations. (**D**), The results of outgroup *f*_*3*_-statistics of the form *f*_*3*_(Target, Ancient Eurasians; Mbuti). Ancient northern East Asians were employed as ancient Eurasian reference populations. (**E**), Maximum-likelihood tree of 50 TB, MG, TK, and Tungusic groups constructed in TreeMix. Modern populations were color-coded according to language classifications and ancient populations were color-coded according to geographical locations. The detailed sample legends are presented in Supplementary fig. S2.

The shared drift measured via the outgroup *f*_*3*_-statistics revealed that target Northwest Chinese populations generally shared more alleles with MG-speaking people from North China, TB and Tungusic-speaking groups (**fig. 1C**). Besides, Northwest Chinese populations also shared more alleles with ancient individuals from the YRB, WLR, Inner Mongolia, Shaanxi, and Nepal (**fig. 1D**). The TreeMix analysis revealed that Salar and Tu populations phylogenetically clustered together with southwestern TB-speaking groups. In contrast, other target populations shared more drift with MG-speaking groups (**fig. 1E**). Zhuoni Tibetan was also phylogenetically close to northwestern TB-speaking groups (**fig. 1E**).

### Characterization of ancestral makeup and admixture landscape

We calculated the admixture *f*_*3*_*-*statistics to explore the potential admixture profile of Northwest Chinese populations. The results of the *f*_*3*_-statistics of the form *f*_*3*_ (Source1, Source2; Target populations) indicated that studied Tibetan, Dongxiang, and Yugur groups were admixed populations and derived from an admixture of East Asian and West Eurasian populations, suggesting that these admixed populations resulted from east-west admixture events (**supplementary tables S1-S4**). Whereas, we did not observe significant negative Z-scores (< −3) in Salar and Tu_groups, i.e.,*f*_*3*_(Source1, Source2;Salar_Xunhua) ≥ 2.025 and *f*_*3*_(Source1, Source2; Tu_Huzhu) ≥ 2.185 (**supplementary tables S5 and S6**), suggested that their relatively isolated genetic background via the unique population history.

Using unsupervised ADMIXTURE analysis, we further dissected the population structure among Northwest Chinese populations and modern and ancient Eurasian populations. When the number of hypothetical ancestral components was assumed to be nine with the lowest cross-validation error (**fig. 2**), the ancestral makeup of Northwest Chinese populations could be explained by four major ancestral components that were maximized in Suila (Tibetan-related, northern East Asian ancestry),BoY(Tai-Kadai-related, southern East Asian ancestry), Mogushan_Xianbei_IA (Northeast Asian ancestry), and French (western Eurasian ancestry), respectively. Tu and Salar populations showed a larger proportion of northern East Asian ancestry, whereas other Northwest Chinese populations harbored a higher proportion of southern East Asian, Northeast Asian, and western Eurasian components. These patterns of population structure were consistent with that unveiled by PCA (**fig. 1B and 1C**). Notably, the fraction of Northeast Asian and western Eurasian ancestries in studied TK-speaking speakers was significantly lower than that in linguistically close reference populations from Xinjiang. Conversely, the fractions of the above two components in Zhuoni Tibetan were higher than that in southwestern TB-speaking groups. Besides, we observed that the Northeast Asian component in studied MG-speaking groups was dramatically lower than that in MG-speaking groups from southern Siberia, while the proportion of the western Eurasian component did not fluctuate significantly among MG-speaking groups.

**Figure 2.**
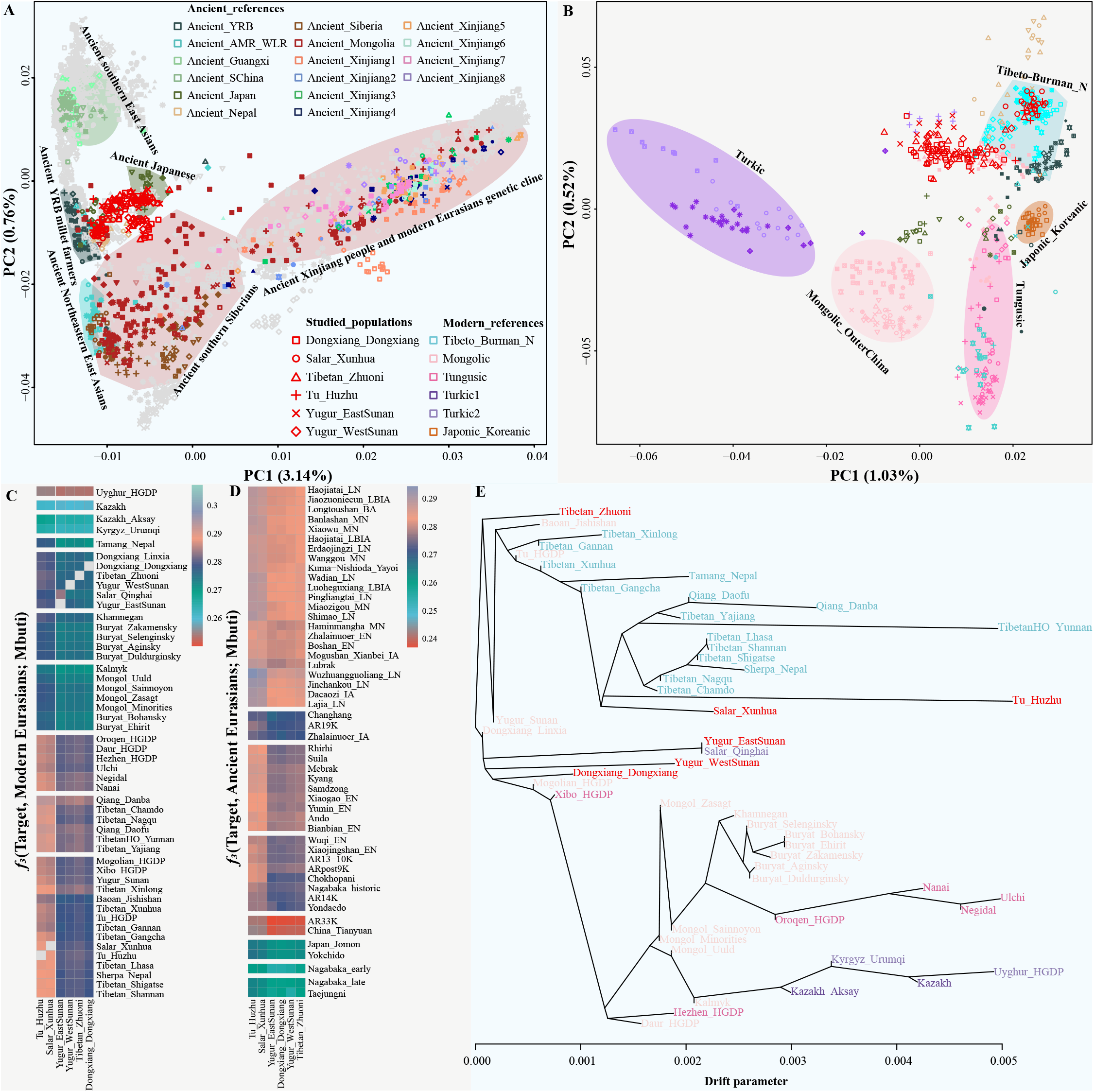
Model-based ADMIXTURE result of target Northwest Chinese populations with modern and ancient Eurasian populations at K = 9. The geographical locations of included modern and ancient East Asian, North Asian, and Southeast Asian populations are displayed in the center of the pie chart. The color-coded legends are consistent with that in PCA. Each ancestral component and corresponding geographical location are labeled. The admixture results and geographical locations of six Northwest Chinese populations are displayed in the surrounding of the center pie chart.

We further used *f*_*4*_*-*statistics to identify ancestry sharing based on shared alleles. We first applied *f*_4_-statistics of the form *f*_4_(Eurasian1, Eurasian2; Target, Mbuti) to evaluate the allele sharing between target populations and Eurasian ancients. We found that studied populations generally shared more alleles with ancient northern East Asians relative to ancient southern East Asians, ancient populations from Xinjiang, Mongolia, and West Eurasia (**supplementary table S7**). Our affinity-*f*_*4*_-based estimates confirmed the observed patterns inferred from the outgroup-*f*_*3*_-statistics. The significantly positive Z-values of *f*_*4*_ (Ancient northern East Asian1, Ancient northern East Asian2; Target, Mbuti) showed that studied populations shared more alleles with post-Middle Neolithic YRB individuals and ancient populations from WLR compared with ancient populations from AR and Nepal (**supplementary table S7**). We also found that target populations shared excess ancestry with Coastal Historic southern East Asians relative to pre-Neolithic AR individuals, and studied Dongxiang, Tibetan, and Yugur populations also shared a closer affinity with Coastal Historic southern East Asians relative to highland Nepal ancients, i.e., *f*_*4*_(China_SEastAsia_Coastal_Historic, Nepal ancients; Dongxiang/Tibetan/Yugur, Mbuti) > 2.98*SE. Furthermore, target populations shared more alleles with Late Neolithic to Historic southern East Asians compared with Bronze Age to Iron Age populations from Mongolia (except Ulaanzuukh and SlabGrave individuals), ancient populations from Xinjiang and Russia (except Ancient Northeast Asians (ANA), such as DevilsCave_N and Boisman_MN), and ancient West Eurasian populations (**supplementary table S7**).

We then calculated *f*_4_(Eurasian1, Target; Eurasian2, Mbuti) to estimate whether additional gene flow contributed to the gene pools of target populations. The significantly negative Z-scores of *f*_*4*_(Nepal ancients, Target; Ancient southern East Asians, Mbuti) showed that Salar and Tu populations shared excess ancestry with Late Neolithic to Historic southern East Asians when compared with ancient populations from Nepal (**supplementary table S8**). We found that, compared with ancient southern East Asians, Salar and Tu populations showed a closer affinity with Middle Bronze Age to Early Iron Age populations from Mongolia (represented by Ulaanzuukh_LBA, SlabGrave_EIA, and Khovsgol _LBA), which possessed major ANA-related genetic profile. Dongxiang, Tibetan, and Yugur populations also shared excess ancestry with these ANA-related populations relative to Early Neolithic southern East Asians. Furthermore, the observed significantly negative Z-scores of *f*_*4*_(Ancient East Asians, Target; Xinjiang ancients/West Eurasians, Mbuti) further showed that West Eurasian representative sources have contributed to the gene pools of target Northwest Chinese populations, i.e., *f*_*4*_(China_SEastAsia_Coastal_EN, Target; Tarim_EMBA1, Mbuti) < -3 (−4.86 < Z < -3.156), *f*_*4*_(China_YR_LN, Target; Basque, Mbuti) < -3 (−13.087 < Z < -3.761), and *f*_*4*_(China_Upper_YR_LN, Target; Russia_MLBA_Sintashta, Mbuti) < -3 (−9.104 < Z < -3.01).

### Genetic differentiation in geographically/linguistically close Northwest Chinese populations

The patterns of genetic relatedness unveiled via PCA, outgroup *f*_*3*_-statistics, and TreeMix analyses showed that genetic differentiations existed among target populations or between studied populations and geographically/linguistically close reference populations. We confirmed the differentiated population structure among Northwest Chinese populations according to our estimates by pairwise qpWave analysis (**supplementary figs. S4 and S5**). Moreover, the pairwise Fst values between Xunhua Salar and Huzhu Tu, or eastern Yugur and western Yugur were close to zero (**supplementary table S9**). Dongxiang population showed the closest relatedness with Zhuoni Tibetan (Fst = 0.0003), followed by Yugur groups (< 0.0011), and Zhuoni Tibetan also showed a relatively high affinity with western Yugur (0.0001) and eastern Yugur (0.0009). We further performed *f*_*4*_*-*statistics of the form *f*_4_(Target1, Target2; Reference populations, Mbuti) to estimate the genetic differentiation between target Northwest Chinese populations. We found that studied Dongxiang, Tibetan, and Yugur populations possessed more West Eurasian ancestry relative to Salar and Tu populations (**supplementary figs. S6 and S7**), and studied Dongxiang and Yugur populations also had more West Eurasian ancestry relative to Zhuoni Tibetan (**supplementary fig. S8A, S8D, and S8E**), which was consistent with the ancestry composition patterns revealed by ADMIXTURE analysis. Generally, Salar and Tu groups showed genetic homogeneity (**supplementary fig. S6C**), Dongxiang and Yugur groups possessed relatively homogeneous genetic backgrounds (**supplementary fig. S8B, S8C, and S8F**).

We next estimated the total and mean lengths of runs of homozygosity (ROH) to identify the genomic footprint of inbreeding. Target Northwest Chinese populations had a similar degree of short ROHs overall, which indicated that they were highly admixed populations and no signs of recent inbreeding was found (**fig. 3A**). We found that the average total length of ROHs in Salar and Tu people was slightly larger than that in other studied individuals (**fig. 3B**). We also observed that, compared with geographically or linguistically close populations (except Uyghur), target populations had fewer mean total length of ROHs (**supplementary fig. S9**). To further investigate the fine-scale genetic structure of target groups, we carried out the haplotype-based identity by descent (IBD) and fineSTRUCTURE analyses. We did not find any strong IBD sharing within studied populations or between these groups and geographically/linguistically close reference populations (**fig. 3C, 3D, 3E, and 3F**), which might be attributed to ancient and extensive interactions of target Northwest Chinese groups with incoming populations. Two major genetic clusters, one including Salars and Tus and the other including Dongxiangs, Yugurs, and Tibetans, and the genetic differentiation between studied groups and linguistically close but more northerly populations were further confirmed by the individual-level bifurcating tree and pairwise coincidence heatmap (**supplementary figs. S10 and S11**).

**Figure 3.**
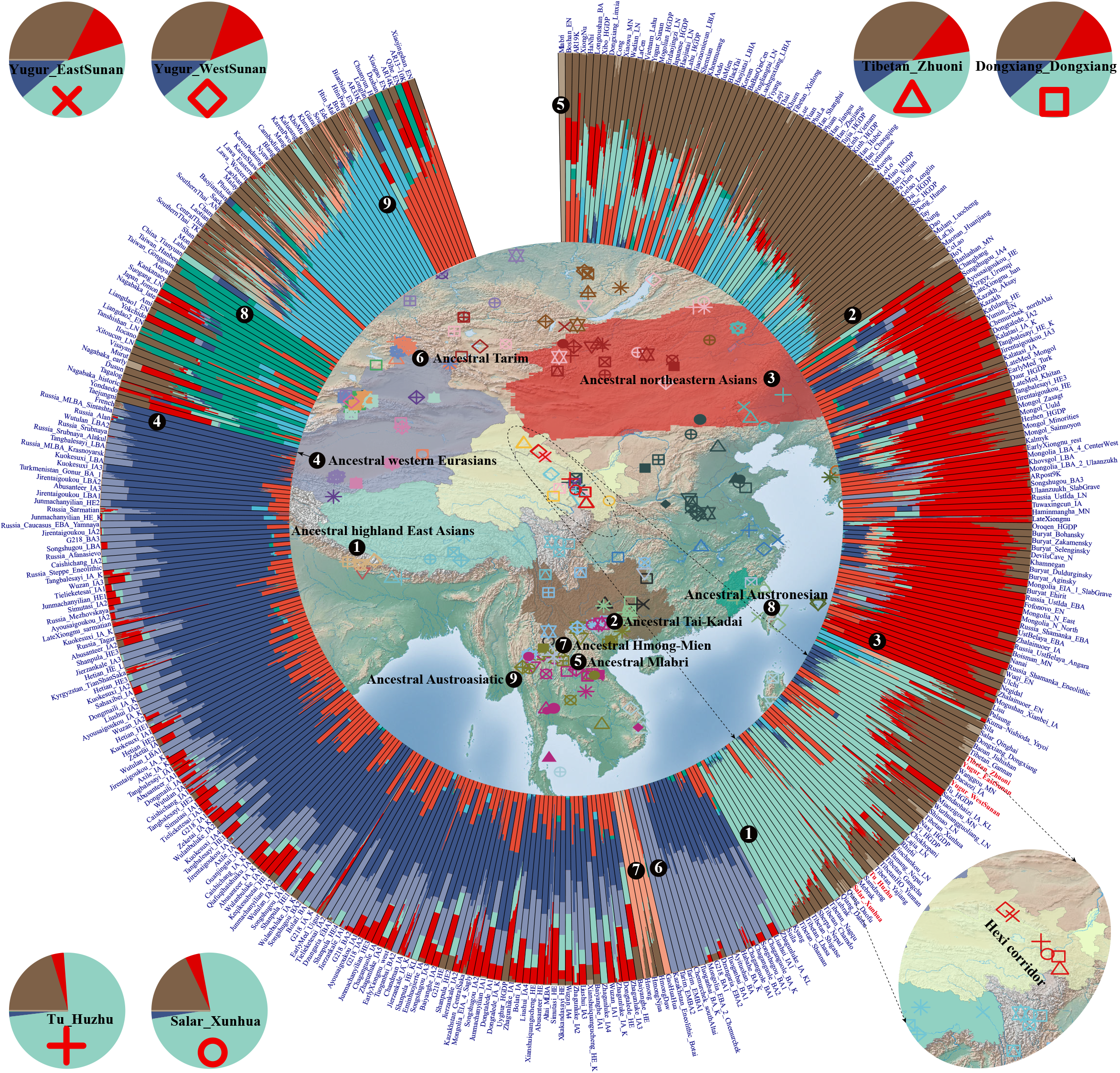
The distribution of runs of homozygosity (ROH) and identity by descent (IBD) sharing between studied Northwest Chinese populations and geographically/linguistically close reference populations based on the whole-genome sequencing dataset. (**A**), The total lengths and numbers of ROH for each studied sample. (**B**), The mean lengths of ROH for each studied population. Network visualizations of the mean length of IBD tracts shared between involved populations, with identified IBD blocks at the overall level (**C**), in the range of 5–10 cM (**D**), 1–5cM (**E**), and less than 1cM (**F**).

### Estimation of admixture proportions and admixture dates

We performed a series of qpAdm analyses to quantify the proportions of different ancestral sources in target populations. Twoway admixture models with Shimao/Upper Xiajiadian/Miaozigou-related populations as one East Eurasian ancestry and Bactria-Margiana Archaeological Complex (BMAC)/Indus periphery/Steppe_EMBA/Steppe_MLBA-derived populations as one West Eurasian ancestry worked out well to explain the formation of studied populations (**fig. 4A**). We found that target populations could be modeled as a mixture of China_Shimao_LN-like (87.2 to 94.4%) and Andronovo-like (Steppe_MLBA-related, 5.6 to 12.8%) sources. A high degree of China_Shimao_LN-related ancestry (84.3 to 94.2%) was further identified in the successfully-fitted qpAdm models, with the rest being derived from Gonur1_BA (BMAC-related, 6.3 to 14.4%) or Dzungaria_EBA1 (6.9 to 15.7%) or Afanasievo (Steppe_EMBA-related, 5.8 to 13.4%) (**fig. 4A**). Target populations could also be modeled as a two-source admixture of China_Miaozigou_MN (77.9 to 89.1%) and Gonur2_BA (Indus periphery-related, 10.9 to 22.1%) or China_WLR_BA (Upper Xiajiadian-related, 74.4 to 87.3%) and Gonur2_BA (12.7 to 25.6%). In several working qpAdm models, target populations (except Salar_Xunhua) could be modeled as a three-source admixture of China_SEastAsia_Coastal_Historic (65.9 to 77.5%), Shamanka_EBA (11.1 to 15%), and Gonur2_BA (8 to 20.7%). We observed that target populations could also be modeled using China_Shimao_LN (77.6 to 89.8%) with additional ancestry from Gonur2_BA (8.1 to 15.7%) and Dzungaria_EBA1 (2.1 to 7.7%) sources (**fig. 4A**). Additionally, models with more potential ancestral sources could depict studied Dongxiang, Tibetan, and Yugur populations as a mixture of Neolithic to Bronze Age East Eurasians (represented by Mongolia_East_N, DevilsCave_N, and Shamanka_Eneolithic), Late Neolithic to Historic southern East Asians, Gonur2_BA, and Steppe-related populations (represented by Dzungaria_EBA2, Tarim_EMBA, and Botai_Eneolithic) (**fig. 4A**). Estimates of admixture dates showed that East/West Eurasian admixture in Dongxiang, Tibetan, and Yugur populations occurred ∼783–841 years ago (27–29 generations), and this admixture in Xunhua Salar occurred approximately 1131 years ago (39 generations). However, we failed to date the eastern-western admixture event in Huzhu Tu due to the small sample size.

**Figure 4.**
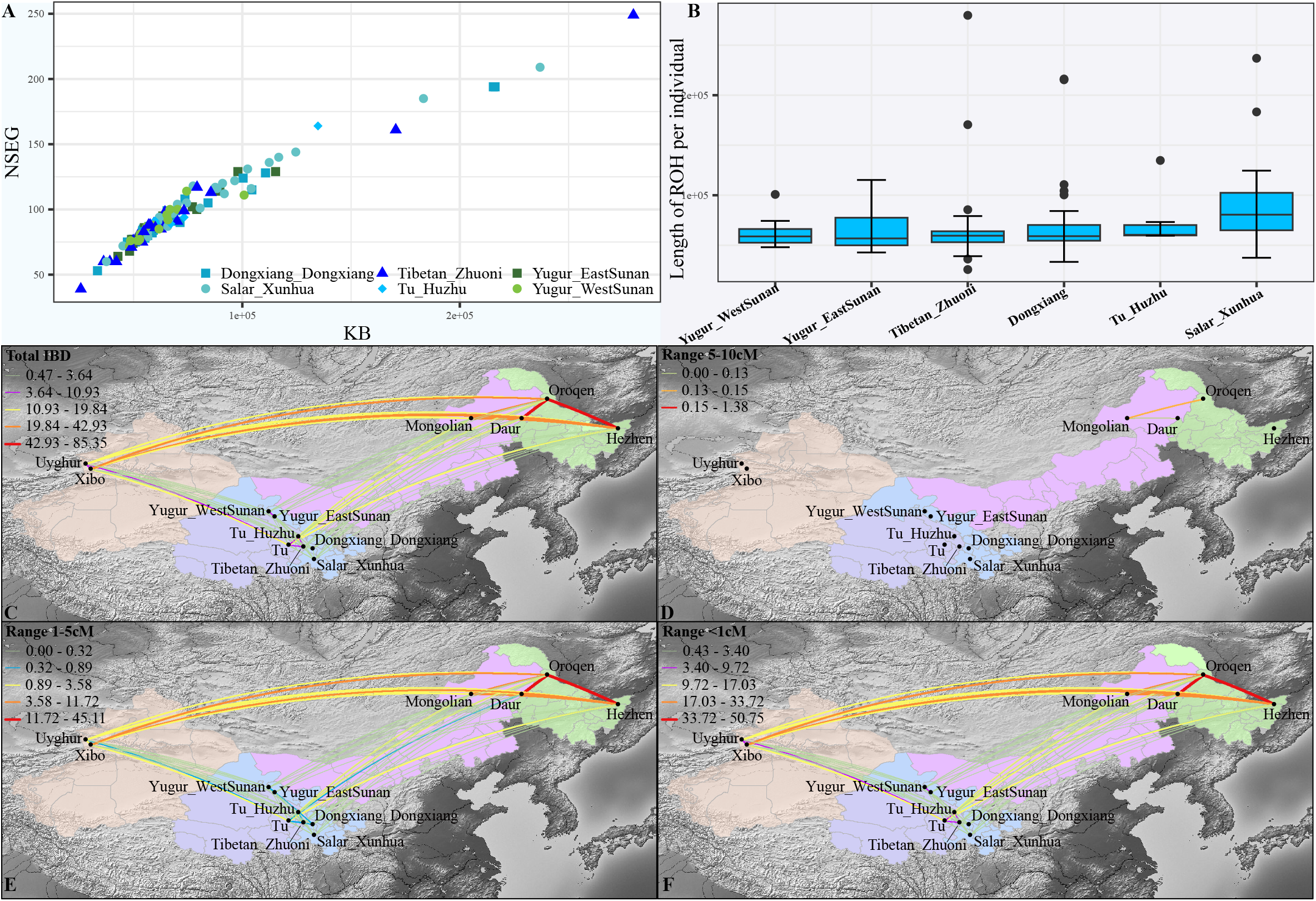
The well-fitted qpAdm admixture models and demographic history of studied Northwest Chinese populations. (**A**), QpAdm admixture models with China_Shimao_LN, China_Miaozigou_MN, China_WLR_BA, Taiwan_Hanben_IA, China_SEastAsia_Coastal_Historic, Russia_Shamanka_EBA, and Russia_DevilsCave_N as the East Eurasian ancestral source and Andronovo, Gonur1_BA, Gonur2_BA, Dzungaria_EBA1, Dzungaria_EBA2, Russia_Afanasievo, and Russia_Caucasus_EBA_Yamnaya as the West Eurasian ancestral source. (**B**), Effective population sizes for six Northwest Chinese populations inferred using SMC++. Seven samples were randomly selected from each population. (**C**), MSMC2 cross-population results for pairs of northern Han Chinese and target Northwest Chinese populations. (**D**), MSMC2 cross-population results for pairs of studied populations.

### Inference of effective population size histories and split times

We applied SMC++ to infer the effective population size (Ne) of Northwest Chinese minority populations, which was estimated by incorporating information from the site frequency spectrum from unphased whole genomes. Target Northwest Chinese populations were inferred to have experienced major growth since the initial peopling of East Asia by anatomically modern humans (**fig. 4B**). We observed that one set of populations, including Dongxiang, Tibetan, and western Yugur populations, and the other one including Salar and Tu populations separately had similar growth patterns, and these groups also experienced some growth after 10 kya. We found that Dongxiang, Tibetan, and western Yugur populations showed an overall larger Ne than other studied populations. Unexpectedly, though two studied Yugur groups exhibited genetic homogeneity, the eastern Yugur had a relatively slow growth rate. We next estimated the time course of population separations using the MSMC2 (multiple sequentially Markovian coalescent) method. The midpoint estimates suggested splits between Northern Han Chinese and Salar_Xunhua ∼14.15 kya; and Tu_Huzhu ∼12.81 kya; and Yugur_WestSunan ∼11.88 kya; and Yugur_EastSunan ∼10.48 kya; and Tibetan_Zhuoni ∼9.73 kya; and Dongxiang_Dongxiang ∼7.68 kya (**fig. 4C**). The estimated split times for 15 population pairs indicated that Northwest Chinese populations began to separate at ∼10.60 kya, and the latest separation occurred between Tibetan and Dongxiang groups ∼7.46 kya (**fig. 4D**).

### Sex-biased admixture

To investigate the unequal contributions from male and female lineages of the ancestral populations to the studied ones, we examined the sex-biased admixture by comparing the admixture results generated based on autosomes, X-chromosome, Y-chromosome, and mtDNA. We found that two-way admixture models with East and West Eurasian ancestral sources could fit the demographic history of target populations. The estimated West Eurasian genetic contribution in target populations was ∼1.1–9.2% for autosomes and ∼2.1–15.2% for X-chromosome (**figs. 5A and 5B**). Uniparental genetic landscape showed that West Eurasian-derived paternal lineages accounted for 33.3–42.4% in studied Dongxiang, Tibetan, and Yugur populations, and corresponding maternal lineages accounted for less than 6.3% (**supplementary table S10, fig. 5C–5E, and supplementary fig. S12**). Notably, we did not observe West Eurasian lineages in Salar and Tu populations, and West Eurasian-typical mtDNA lineages were also not detected in western Yugur and Tibetan populations (**fig. 5D and 5E**). Generally, the different contributions of West Eurasian immigrants to the paternal and maternal lineages and the significant difference in admixture proportions between autosomes and X-chromosome indicated that the admixture history of Dongxiang, Tibetan, and Yugur populations was sex-biased with an excess of East Asian females and West Eurasian males.

**Figure 5.**
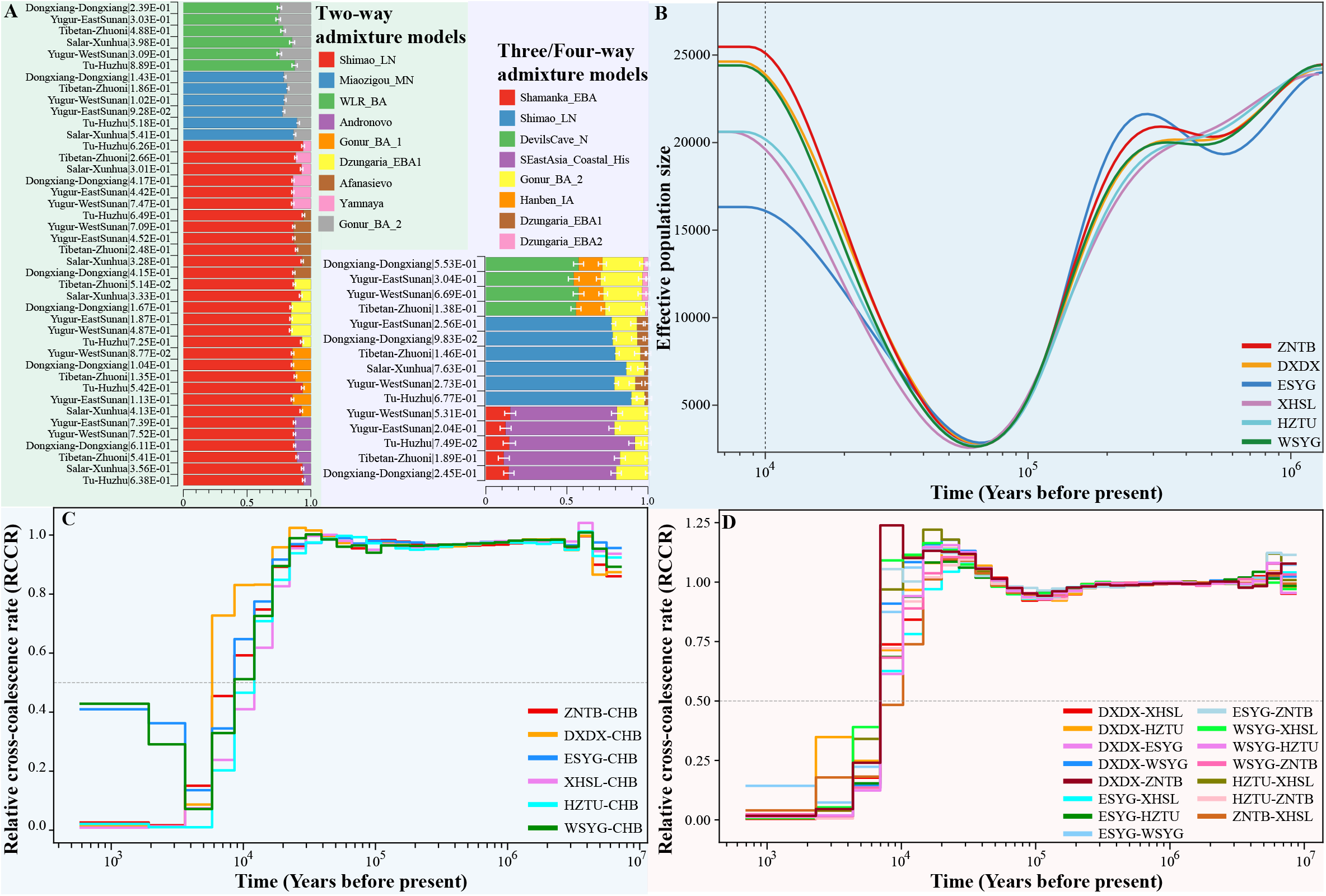
Sex-biased admixture history of target Northwest Chinese populations. (**A**), The ancestral makeup of six Northwest Chinese populations estimated based on autosomal genetic variants from the whole-genome sequencing dataset. (**B**), The ancestral makeup of six Northwest Chinese populations estimated based on X-chromosomal genetic variants from the whole-genome sequencing dataset. (**C**), The proportions of East and West Eurasian ancestries in target Northwest Chinese populations estimated based on markers on different chromosomes. The haplogroup distribution of paternal lineages (**D**) and maternal lineages (**E**).

### Identification of signatures of natural selection

We calculated each group’s population branch statistic (PBS) to detect the natural selection signatures in studied populations. We used Han Chinese and European populations from the HGDP as the second and third reference populations, which allowed us to detect the genes that were likely under selection in target Northwest Chinese populations but not in Han Chinese. We found that the Salar/Tu population had more significant selection signals than the other three populations, and Northwest Chinese populations with a higher West Eurasian ancestral component tended to share more adaptive signals. We observed that *PRIM2, NBPFs 9/20/25P*, and *PDE4DIP* showed evident signatures of natural selection in all studied populations (**fig. 6A–6D**). *PRIM2* plays a crucial role in the replication of DNA, which was identified by a large-scale genome-wide association study to be associated with coronary artery disease (Van Der Harst and Verweij 2018). *NBPFs 9/20/25P* are members of the neuroblastoma breakpoint family, which have been implicated in several developmental and neurogenetic diseases. Moreover, *NBPF8*, also a neuroblastoma breakpoint family member, was identified as a significant selection signal in Dongxiang, Tibetan, and Yugur populations (**fig. 6A–6C**). Recent genetic findings provided compelling evidence for an association between the *PDE4DIP* mutations and heart block, atrial fibrillation, and heart failure (Abou Ziki, et al. 2020; Abou Ziki, et al. 2021; Mani 2022). In addition, the *FRG1* gene was identified as a significant selection signal in Dongxiang, Tibetan, and Yugur populations, which is associated with facioscapulohumeral muscular dystrophy (**fig. 6A–6C**). *NOTCH2* gene, which plays a role in vascular, renal, and hepatic development, showed relatively high PBS values in Tibetan and Yugur populations (**fig. 6B and 6C**). We found that the *TSNAX-DISC1* gene associated with risk for psychiatric illness, most notably schizophrenia was an extremely significant selection signal in Salar/Tu group (**fig. 6D**). Unexpectedly, the *EPAS1* gene, which is associated with high-altitude adaptation, was also identified as an adaptive signal in Salar/Tu group, and *EPAS1* variants in Dongxiang and Yugur populations showed lower PBS values (0.10–0.16) than that in Salar/Tu population (0.11–0.43) (**fig. 6A, 6C, and 6D**). Allele frequency spectrum of the selected allels showed significant difference between East Asians and other continental populations, such as the top selected loci in the TSNAX-DISC1 (**fig. 6E, 6F**). In general, Dongxiang, Tibetan, and Yugur populations shared more natural selection signals with each other than with Salar/Tu group (**fig. 6G**).

**Figure 6.**
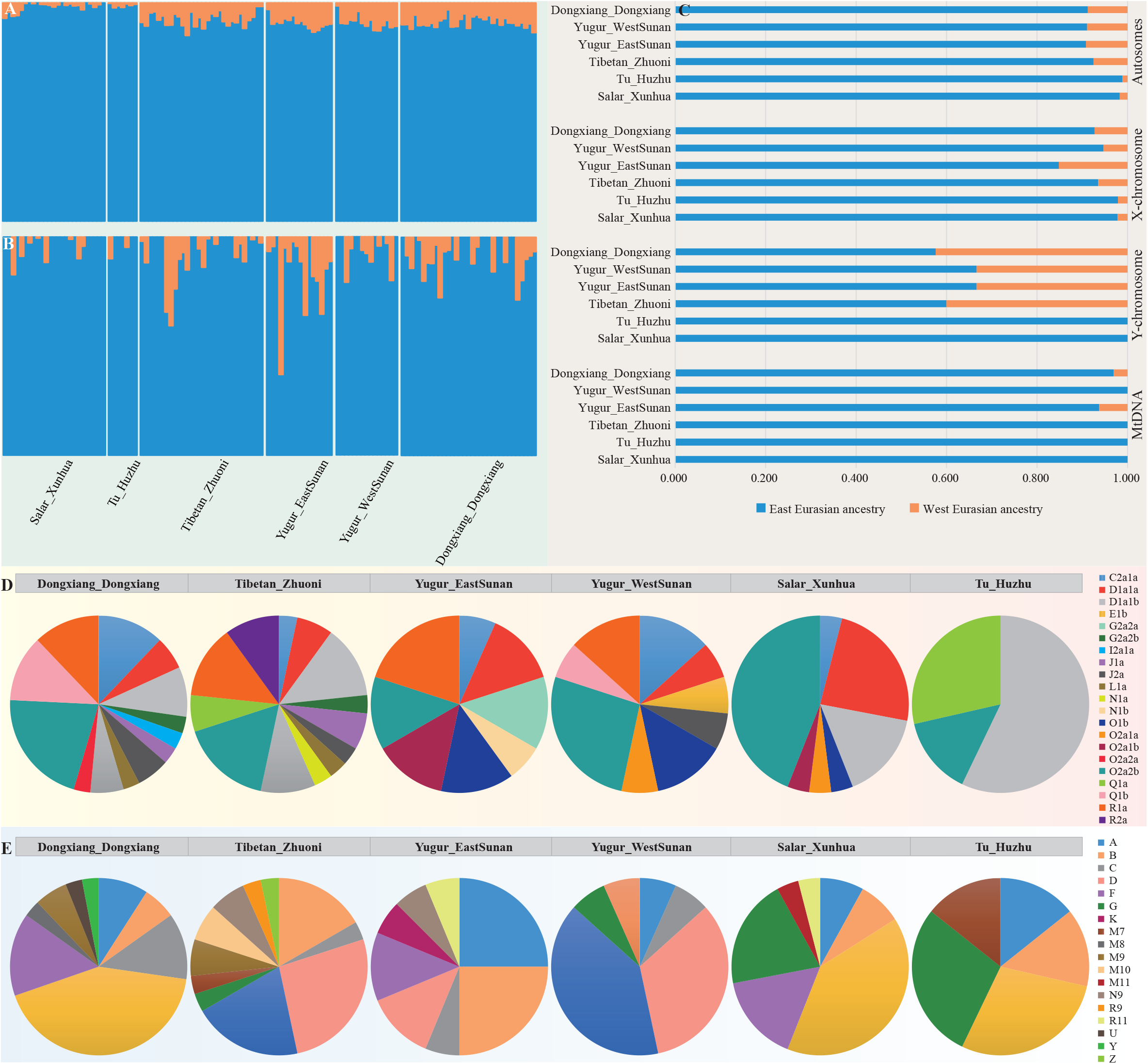
The signals of natural selection in studied Northwest Chinese populations screened via the PBS approach and using Han Chinese and European populations as the close and distant reference populations. The natural selection signals identified in Dongxiang Dongxiang (**A**), Zhuoni Tibetan (**B**), eastern and western Yugur populations (**C**), and Xunhua Salar/Huzhu Tu (**D**). The candidate genes ranked at top 100 and involved at least three SNPs are labeled. Due to the overlap of positions between candidate genes, only the first occurrence is labeled. (**E∼F**) Allele frequency of two selected loci (rs6655954 and rs78658433) in TSNAX-DISC1 among 380 worldwide populations from the 10K_CPGDP, HGDP and Oceacina genomic sources. (**G**), The Circos and upset plots showed overlaps among the input gene lists. On the outside, each arc represents the identity of each gene list. On the inside, each arc represents a gene list. The dark orange color represents the genes shared by multiple lists and the light orange color represents genes unique to that gene list. Purple lines link the same genes that are shared by multiple gene lists. The greater the number of purple links and the longer the dark orange arcs imply greater overlap among the input gene lists. (**H**), Heatmap of enriched terms across input gene lists, colored by p-values.

Given that target Northwest Chinese populations could be modeled as a mixture of East (represented by Ataya, Paiwan, and other populations in southern East Asia) and West Eurasian ancestral sources, we further performed PBS analysis by employing southern East Asian and European populations as the close and distant reference populations. The *PRIM2* gene was again identified as a significant selection signal in all studied populations (**supplementary fig. S13**). Moreover, we found that *DDAH1*, which is associated with cardiovascular system disease and pulmonary hypertension, was a shared selection signal in Dongxiang, Tibetan, and Yugur populations (**supplementary fig. S13A–S13C**). The *GALNT2* gene, which may influence triglyceride levels and may be associated with Type 2 diabetes as well as several types of cancer, was identified as a significant selection signal in Zhuoni Tibetan (**supplementary fig. S13B**). *MLIP* gene, which is predicted to be involved in the negative regulation of cardiac muscle hypertrophy in response to stress, exhibited high PBS values in Salar/Tu population (**supplementary fig. S13D**). We found that *KHDRBS2*, which plays a role in regulating alternative splicing of Neurexin 1 in the laminin G-like domain 6 involved in neurexin selective targeting to postsynaptic partners, showed significant signals of selection in Zhuoni Tibetan (**supplementary fig. S13B**). The *SIPA1L2* gene was reported to be associated with Parkinson’s disease in East Asians (Chen, et al. 2016; Zou, et al. 2018) and was identified as an adaptive signal in Salar/Tu population. The *EHBP1* gene, associated with an aggressive form of prostate cancer, was also identified as a signal of strong selection in Salar/Tu population. Additionally, among the top 100 SNPs, rs7589861 and rs13420272 located in the *EPAS1* gene showed high genetic differentiation between Salar/Tu and southern East Asian ancestral populations (**supplementary fig. S13D**).

We then conducted an enrichment analysis to confirm the biological processes of identified candidate genes when using Han Chinese as the reference population. We found that these candidate genes are associated with one WikiPathways and five GO (gene ontology) biological pathways (**fig. 6H**). The VEGFA-VEGFR2 signaling pathway (WP3888) is essential for modulating angiogenesis by regulating endothelial permeability and the proliferation, survival, sprouting, and migration of endothelial cells (Lohela, et al. 2009; Claesson-Welsh and Welsh 2013). These five GO pathways are respectively associated with positive regulation of GTPase activity (GO: 0043547), regulation of the frequency, rate, or extent of small GTPase-mediated signal transduction (GO: 0051056), neuron projection morphogenesis (GO: 0048812), cerebral cortex radial glia-guided migration (GO: 0021801), and regulation of cell growth (GO: 0001558).

## DISCUSSION

Discovering and understanding human genetic variations as well as characterizing the full landscape of human genomic diversity of under-represented populations in large-scale international genomic reference databases was necessary for personal health management and more accurate disease risk prediction in precision medicine. In the present study, we have generated and analyzed whole-genome sequencing data from six ethnolinguistic groups in Northwest China. In this set of 126 genomes, we identified approximately 16.4 million SNPs and 1.9 million InDels, including hundreds of thousands of previously undocumented variants. We point to the importance of sequencing ethnolinguistically diverse East Asians to understand population migration and admixture history as well as the determinants of health disparity of Northwest Chinese populations.

With this unprecedented data, we comprehensively unraveled the population structure and admixture history of target Northwest Chinese populations. We found that Northwest Chinese populations generally showed a close affinity with TB, MG, Tungusic groups, and ancient populations from YRB, WLR, Nepal, Japan, and Mongolia (**fig. 1A, supplementary figs. S1A, S2, and S3**). Though these studied populations clustered together with aforementioned modern and ancient northern East Asians, we found that they were genetically differentiated and could be divided into two groups, with one including Salar and Tu people and the other including Dongxiang, Tibetan, and Yugur people. Specifically, the patterns of population structure revealed by PCA, outgroup *f*_*3*_-statistics, TreeMix, and pairwise qpWave analyses showed that Salar and Tu populations shared more alleles with southwestern TB groups and ancient individuals from Nepal, and other target populations shared more alleles with MG and Tungusic groups as well as ancient populations from southern Siberia (**figs. 1B, 1E, supplementary figs. S1B, S4, and S5**). Moreover, Zhuoni Tibetan also shared more alleles with northwestern TB groups. In short, the patterns of genetic structure were not consistent with linguistic classifications, especially for studied TK groups.

Previous studies based on genome-wide SNP data revealed that Dongxiang, Bonan, Yugur, Salar, and Hui populations in Gansu province, Northwest China derived 9–15% of their ancestry from West Eurasians (Ma, Chen, et al. 2021), and Gansu Tibetans presented a low level of West Eurasian admixture (Yao, et al. 2017). Our observations confirmed the eastern-western admixture in the newly-studied Northwest Chinese populations. We identified four major ancestral components in target populations, namely northern East Asian, southern East Asian, Northeast Asian, and West Eurasian ancestries (**fig. 2**). The results of *f*_4_(Eurasian1, Eurasian2; Target, Mbuti) and *f*_4_(Eurasian1, Target; Eurasian2, Mbuti) further demonstrated that target populations shared excess ancestry with these potential ancestral sources to a varying degree. We found that twoway admixture models with East and West Eurasian ancestries could well explain the admixture history of target populations. Also, models with more potential ancestral sources also worked well to explain most studied Northwest Chinese populations’ formation. However, there were no available threeway admixture models that showed a good fit for Xunhua Salar, and no available fourway admixture models for Salar and Tu populations. A recent study estimated that Gansu Salar derived ∼11.8% of their ancestry from West Eurasians (represented by Andronovo samples), which was higher than our estimation (∼6.5%). Generally, target populations derived their ancestry primarily from East Asians. Though Dongxiang and Salar people practice Islam, our findings supported that the cultural diffusion model played a pivotal role in the formation of these two populations. Besides that, we found that Dongxiang, Tibetan, and Yugur populations possessed a higher proportion of West Eurasian ancestry relative to Salar and Tu populations (**fig. 2**). This finding was confirmed by *f*_4_(Target1, Target2; Eurasians, Mbuti) and the working qpAdm models (**fig. 5 and supplementary figs. S6–S8**). Generally, the genetic differentiation among target Northwest Chinese populations might be attributed to the difference in the proportion of West Eurasian ancestry.

The estimation of Ne reflected differentiated growth patterns among studied populations. That is, the growth rates of Dongxiang, Tibetan, and western Yugur groups were higher than that in other studied populations. The differentiated growth patterns between the two Yugur groups were inconsistent with their homogeneous genetic background revealed by allele-based analyses. We found that the split times between Northern Han Chinese and target populations were dated at ∼7.68–14.15 kya, and the separation between studied populations occurred at ∼7.46–10.60 kya. To a certain extent, the Holocene climatic change and the development of Neolithic culture (especially in agriculture) facilitated human migration and separation (Li, et al. 2015; Zhao 2020). Specifically, the divergence between Tibetan and Han Chinese was estimated to occur ∼9.73 kya, which was consistent with the estimation based on deeply sequenced genomes of 38 Tibetan highlanders and 39 Han Chinese lowlanders (Lu, et al. 2016) but higher than the split time measured based on Bayesian phylogenetic analysis of Sino-Tibetan language family (Zhang, et al. 2019). We estimated that the eastern-western admixture occurred ∼783–1131 years ago, roughly dating back to Tang Dynasty, the Five Dynasties and Ten Kingdoms period, and Song Dynasty. According to historical records, the ancient Silk Road was reopened by the Tang Empire in AD 639, and intensive contact between East and West Eurasian populations was common during the early Tang Dynasty, indicating that the admixture dates might be underestimated. The admixture history of target Northwest Chinese populations could be more complex than our estimated models.

Sex-biased demographic events have played a vital role in shaping human genetic diversity. Xiong et al. analyzed the uniparental lineages in Heishuiguo samples and found that male-dominated migration by YRB immigrants might significantly affect the subsistence strategy transition of historical populations along the Hexi Corridor (Xiong, et al. 2022). Ma et al. identified considerable sex-biased admixture, involving an excess of western males and eastern females, in the gene pool of Ningxia Huis (Ma, Yang, et al. 2021). Based on genome-wide SNP data, the male-dominant sex-biased admixture events were also identified in Dongxiang, Bonan, Yugur, and Salar populations (Ma, Chen, et al. 2021). We identified sex-biased admixture, with more East Eurasian females and West Eurasian males, in the demographic history of our studied Dongxiang, Tibetan, and Yugur groups. According to historical records, traders, soldiers, emissaries, and scholars who migrated from Central Asia, Arabia, Persia, and other western regions into China were mostly males. They transmitted West Eurasian-typical paternal lineages to some present-day Northwest Chinese individuals. However, we did not observe sex-biased gene flow in Salar and Tu people, which was not consistent with previous findings (Wang, et al. 2003; Yao, et al. 2016), indicating that genetic differentiation may exist in geographically diverse Salars. Moreover, the sampling bias and limited sample size might result in underestimating West Eurasian-related paternal lineages.

We searched for signatures of natural selection in studied Northwest Chinese populations by scanning functional SNPs and corresponding genes using the PBS approach. We identified five genes (*PRIM2, NBPFs 9/20/25P*, and *PDE4DIP*) that showed strong signatures of natural selection in all studied populations. *PRIM2* and *PDE4DIP* genes were reported to be associated with cardiovascular diseases (Van Der Harst and Verweij 2018; Abou Ziki, et al. 2020; Abou Ziki, et al. 2021; Mani 2022). Furthermore, *NOTCH2*, related to vascular, renal, and hepatic development, was identified as a significant selection signal in Tibetan and Yugur populations; *DDAH1*, associated with cardiovascular system disease and pulmonary hypertension, was also identified as an adaptive signal in Dongxiang, Tibetan, and Yugur populations; *GALNT2*, which plays a role in triglyceride levels and Type 2 diabetes, showed high PBS values in Zhuoni Tibetan; and *MLIP* gene, which is associated with negative regulation of cardiac muscle hypertrophy in response to stress, was identified as a significant selection signal in Salar/Tu population. It is generally known that Northwest China experiences cold and dry winters and hot summers, making calories and salt supplementation more important. A strong flavor is very important for northwesterners, and northern dishes are oilier and richer in meat. Hence, the incidence of cardiovascular diseases is relatively high in Northwest Chinese populations (Li, Wu, et al. 2020), which could lead to the detection of the above selection signatures in studied populations. We found that *NBPFs 8/9/20/25P*, which play a role in a number of developmental and neurogenetic diseases, showed significant signals of natural selection in studied populations. *KHDRBS2*, which is involved in neurexin selective targeting to postsynaptic partners, showed high PBS values in Zhuoni Tibetan. Besides, we found that *TSNAX-DISC1* and *SIPA1L2* (Nakata, et al. 2009; Chen, et al. 2016; Zou, et al. 2018), associated with risk for psychiatric illness, were identified as significant selection signals in the Salar/Tu population. Gansu and Qinghai are two of the least developed provinces in China, people living in rural areas are disproportionately affected in terms of access to health care, medicine accessibility, and so on (Wang, et al. 2009; Yang, et al. 2022), making the health threats to people in underdeveloped regions more severe than urban residents, which might result in the identification of development/neurogenesis-related selection signatures. In the previous study, we observed that *NBPFs 9/10* showed selection signals (using northern Han as a reference population) in Hmong-Mien groups living in Xuyong, Sichuan (Liu, Xie, et al. 2021). Xuyong was a national-level poverty-stricken county before 2020. The harsh living environment of local residents could lead to the detection of development/neurogenesis-related genes. In addition, the *EPAS1* gene was identified as a signature of natural selection in studied populations (except Zhuoni Tibetan). The sampled Northwest Chinese individuals live in areas with high altitudes (northeastern Qinghai-Tibet Plateau), which might contribute to detecting hypoxia-related signatures.

Considering the importance of Northwest China in historical trans-Eurasian cultural and human communication and the unique history of Northwest minorities, the newly-generated whole-genome sequencing data is of great significance for exploring the genetic diversity of East Asian populations and provides reference data for understanding the genetic basis of local adaptation and the genetic architecture of many common and rare diseases. Recruiting more ethnolinguistically and geographically diverse Northwest Chinese populations, conducting high-depth sequencing (≥ 30X) or single-molecule third-generation sequencing, and developing more sophisticated algorithms will promote our understanding of population origin, separation, admixture, adaptation, and introgression from archaic individuals of East Asians.

## CONCLUSION

In this study, we generated whole-genome sequencing data from six MG/TK/TB ethnolinguistic groups from Northwest China and characterized the studied populations’ population structure, admixture history, and local adaptation. Our observations revealed that target Northwest minorities showed excess allele sharing with reference MG, TB, and Tungusic groups from North China as well as Middle Neolithic to Iron Age northern East Asians. In addition, we found that northwestern Chinese populations showed substantial ethnicity/language-correlated genetic differentiation. That is, Dongxiang, Tibetan, and Yugur people had a higher proportion of West Eurasian ancestry. As descendants of residents along the ancient Silk Road, these groups experienced extensive genetic exchanges with incoming populations. The eastern-western admixture was estimated to occur around 783–1131 years ago. We identified considerable sex-biased admixture with more eastern females and western males contributing to the gene pool of studied Dongxiang, Tibetan, and Yugur populations. Moreover, the signatures of natural selection associated with cold arid and high-altitude adaptations were identified in studied Northwest Chinese populations.

## MATERIALS AND METHODS

### Subject details

Saliva samples of 126 unrelated individuals were collected from Northwest China (Yugurs from the western part of Sunan Yugur Autonomous County of Gansu, n=15; Yugurs from the eastern part of Sunan Yugur Autonomous County of Gansu, n=16; Dongxiangs from Dongxiang Autonomous County of Gansu, n=33; Tus from Huzhu Tu Autonomous County of Qinghai, n=7; Salars from Xunhua Salar Autonomous County of Qinghai, n=25; and Tibetans from Zhuoni County in the Gannan Tibetan Autonomous Prefecture of Gansu, n=30). The individuals enrolled in this study were randomly chosen, and informed consent was obtained from all participants. Each participant was the offspring of non-consanguineous marriages of indigenous people of the same nationality within at least three generations. This study was approved by the Medical Ethics Committee of Sichuan University and all procedures were performed in accordance with the Helsinki Declaration of 2013 (Jama 2013).

### DNA extraction and whole-genome sequencing

Genomic DNA was extracted using the QIAamp DNA Mini Kit (QIAGEN, Germany) and quantified using the Qubit dsDNA HS Assay Kit on a Qubit 3.0 fluorometer (Thermo Fisher Scientific) according to the manufacturer’s protocols. Paired-end 150bp whole-genome sequencing was performed on the DNBSEQ-T7 platform (MGI, Shenzhen, China) according to provided protocols with standard library preparation. The target sequencing depth was 10X for all samples.

### Quality control and genotype calling

Each sample was run on a unique lane with at least 30 GB of data that had passed the first filtering step, and each lane was quality controlled after the run to ensure that 80% of the bases achieved at least a quality score of 30. We used Sentieon Genomics software v.201808.01 to perform variant calling following the bioinformatics pipeline for DNA analysis recommended in the Broad institute best practices described in https://www.broadinstitute.org/gatk/guide/best-practices. In brief, we mapped the raw sequencing reads to the human reference genome GRCh37 (human_g1k_v37) using BWA v.0.7.13 (Li and Durbin 2009), and duplicate reads were removed using the Dedup algorithm implemented in Sentieon Genomics. We performed germline joint variant calling analysis using the Sentieon GVCFtyper algorithm. The GATK VariantEval tool was used to calculate various quality metrics of the identified variants.

### Dataset arrangement

Variants from the HGDP (Bergstrom, et al. 2020a) and Oceanian populations (Choin, et al. 2021) were incorporated into this study as the whole-genome sequencing dataset. Before merging these three datasets, multi-allelic variants were removed. Additionally, before merging with data from this study and Oceanian populations, the genome positions of the HGDP were lifted over from GRCh38 to GRCh37. The resulting whole-genome sequencing dataset contained 1,372 samples and 86,629,844 variants. This dataset was mainly used for analyses that need a high marker density, such as inferring the Ne and population split times.

We then merged our dataset with two sets of worldwide genotype panels, one based on the HO dataset (∼600,000 SNPs) and the other on the 1240K dataset (including all SNPs included in the HO dataset) retrieved from the Allen Ancient DNA Resource (AADR, https://reich.hms.harvard.edu/allen-ancient-dna-resource-aadr-downloadable-genotypes-present-day-and-ancient-dna-data). We used PLINK v.1.9 (Chang, et al. 2015) to perform quality control with the following parameters (mind: 0.01, geno: 0.01, maf 0.01, and hwe 10^−6^), and then we used PLINK and King (Manichaikul, et al. 2010) to estimate relatedness coefficients among each pair of individuals from the same population. At last, a merged HO dataset with 13,181 individuals and 593,050 SNPs and a merged 1240K dataset with 10,487 individuals and 1,135,264 SNPs have been generated.

### Principal component analysis and Fst calculation

We used the smartpca package implemented in EIGENSOFT (Patterson, et al. 2012) to conduct PCA clustering by projecting ancient individuals onto present-day ancestries (numoutlieriter: 0 and lsqproject: YES). We first conducted a Eurasian PCA analysis using the whole-genome sequencing dataset and we only employed bi-allelic autosomal SNPs with MAF > 0.05. To explore the fine-scale population structure at broad temporal and spatial scales, we subsequently performed a series of PCAs by gradually removing “outliers” on the first two PCs and re-analyzed the merged HO dataset based on the same set of SNPs. Moreover, we calculated pairwise Fst values between studied populations and geographically/linguistically close reference populations using PLINK v.1.9. To reduce the deviation of Fst values caused by sampling bias, we randomly extracted ten samples from each population, and all samples were included when the sample size of a population was less than ten.

### Model-based ADMIXTURE analysis

We pruned SNPs in strong linkage disequilibrium following the parameters of --indep-pairwise 200 25 0.4 using PLINK v.1.9 (Chang, et al. 2015). We then performed the unsupervised ADMIXTURE analyses using ADMIXTURE v.1.3.0 (Alexander, et al. 2009), with the number of ancestral components K from 2 to 20 and 10-fold cross-validation (−-cv = 10).

### F-statistics

We estimated the shared genetic drift between target populations and Eurasian references using qp3Pop package in ADMIXTOOLS (Patterson, et al. 2012) via outgroup *f*_*3*_-statistics of the form *f*_*3*_(Reference, Target; Mbuti). We also conducted admixture *f*_*3*_-statistics of the form *f*_*3*_(Source1, Source2; Target) to explore the potential admixture signatures of the focused populations, in which significantly negative *f*_*3*_-values with Z-scores less than -3 denoted the admixture evidence involving two predefined ancestry surrogates. Furthermore, we made inferences about the admixture of target populations using qpDstat package in ADMIXTOOLS (Patterson, et al. 2012) with *f*_*4*_ model (*f*_*4*_Mode: YES).

### QpWave/qpAdm analysis

We conducted pairwise qpWave analysis to test whether focused population pairs (left populations) formed one clade compared to a set of outgroups (right populations) via the rank tests. We then used qpAdm package from ADMIXTOOLS (Patterson, et al. 2012) to model the studied populations as an admixed result of groups related to different ancestry surrogates for the true source populations and estimate corresponding admixture coefficients. Two additional parameters “allsnps: YES and details: YES” were used in the qpAdm analysis. We used a basic set of nine outgroups: Mbuti, Russia_Ust_Ishim_HG, Russia_Kostenki14, Belgium_UP_GoyetQ116_1, Italy_North_Villabruna_HG, Israel_Natufian, Atayal, Mixe, and Onge. Also, Turkmenistan_Gonur_BA_1, Russia_MLBA_Sintashta, Russia_Srubnaya, Russia_Caucasus_EBA_Yamnaya, Russia_Afanasievo, Tarim_EMBA1, Tarim_EMBA2, Dzungaria_EBA1, Dzungaria_EBA2, China_NEastAsia_Coastal_EN, China_NEastAsia_Inland_EN, China_YR_MN, China_SEastAsia_Island_EN, China_SEastAsia_Coastal_EN, Mongolia_East_N, Russia_DevilsCave_N, Russia_MN_Boisman, and Russia_Shamanka_Eneolithic were employed as additional outgroups in turn.

### Admixture graph construction

We built the maximum likelihood tree with migration edges to recapitulate population splits and mixtures using TreeMix (Pickrell and Pritchard 2012).

### Admixture time estimation

We used ALDER (Loh, et al. 2013) to date the admixture events with predefined ancestral donor populations to our focused recipient populations, and two additional parameters of jackknife:YES and mindis: 0.005 were used. We only extracted the simple characteristics of a single pulse-like admixture.

### Runs of Homozygosity

We applied PLINK v.1.9 (Chang, et al. 2015) to identify the individual-level ROH. We removed SNPs with a missing genotype rate > 0.05 and MAF < 0.05, then we estimated ROH using a window of 500 kb and allowing one heterozygous SNP per window.

### Identity by descent and FineSTRUCTURE analyses

We used Segmented HAPlotype Estimation & Imputation Tool (SHAPEIT v.2.0) (Delaneau, et al. 2011) for the estimation of haplotypes based on the recommended human genetic maps (Browning and Browning 2013). Subsequently, we used Refined IBD (Browning and Browning 2013) for detecting shared IBD segments within and between individuals or populations (Browning and Browning 2011). We used fineSTRUCTURE v.4.0 to calculate the coancestry matrix and explore the fine-scale population structure based on the haplotype data with default parameters.

### Effective population size and divergence time

We used the SMC++ program (Terhorst, et al. 2017) to estimate the effective population size histories for each of the studied populations separately. We ran SMC++ by assuming a mutation rate of 1.25 × 10^− 8^ per base per generation and a generation time of 29 years. The two samples with the highest sequencing coverage were specified as the distinguished lineages for each population. To reduce the bias caused by different sample sizes, we chose seven samples from each sampled population, which was the minimum sample size of the studied populations.

We applied MSMC2 (Schiffels and Wang 2020) to infer the demographic history and population structure through time. Two samples were randomly selected from each target population. We first used bamCaller.py from the MSMC-tools package to generate sample-specific VCF and mask files. We then statistically phased the VCFs against the 1000 Genomes Phase III reference panel using SHAPEIT v.2.0. For generating multihetsep files for two-phased diploid individuals on each chromosome, we adopted the mappability masks for the human reference genome hs37d5 and used generate_multihetsep.py script from MSMC-tools package to merge VCF and mask files together. Subsequently, MSMC2 was run three times independently for estimating coalescence rates within population1/population2 and across populations. We also used combineCrossCoal.py in the MSMC-tools repository to create a single joint output file with all three estimates obtained from the three MSMC2 runs above. We calculated the absolute time estimation by assuming a slow mutation rate of 1.25 × 10^−8^ per base per generation for a generation time of 29 years. We then computed the relative cross coalescence rate (rCCR) and estimated the divergence time between two populations at which the rCCR hit 0.5.

### Identification of natural selection signals

We applied the PBS approach (Yi, et al. 2010) to detect the signals of natural selection in our studied populations. We only kept SNPs with a more than 95% genotyping rate for PBS calculation. We used Han Chinese or southern East Asian and European populations from the whole-genome sequencing dataset as the close and distant reference populations, respectively. To reduce the bias caused by the small sample size, we combined Salar and Tu populations into one group and the eastern Yugur and western Yugur populations into another group due to their similar genetic backgrounds. We focused on the SNPs showing extreme PBS values (top 100) in target populations as strong candidates for the genetic basis of adaptive evolution. The candidate genes involving at least three functional SNPs were identified as extremely significant selection signals in studied populations.

### Uniparental haplogroup assignment

The Y-chromosomal haplogroups were classified using Y-LineageTracker (Chen, et al. 2021) based on known Y-SNPs from ISOGG Y-DNA Haplogroup Tree 2019-2020. The maternal haplogroups were classified using HaploGrep2 (Weissensteiner, et al. 2016) based on PhyloTree17.

## Acknowledgments

This work was supported by grants from the National Natural Science Foundation of China (82202078). We thank Prof. Etienne Patin and Prof. Lluis Quintana-Murci from the Human Evolutionary Genetics Unit of Institut Pasteur for sharing the high-coverage genomes of 317 individuals from the Pacific region.

## Disclosure of potential conflict of interest

The authors declare that they have no conflict of interest.

## Notes

### Competing Interest Statement

The authors have declared no competing interest.

